# MYBL2 maintains stemness and promotes theta-mediated end joining in triple negative breast cancer

**DOI:** 10.1101/2025.04.15.648875

**Authors:** Rachel Bayley, Anna Munsey, Suman Ahmed, Ciara Ward, Amber Stiby, Bohdana-Myroslava Briantseva, Kirendeep Jawanda, Abeer M Shaaban, Marco Saponaro, Graeme C M Smith, Robert B Clarke, Clare Davies, Paloma Garcia

## Abstract

Triple negative breast cancer (TNBC) has a poor prognosis due to limited treatment options and high metastasis risk. Breast cancer (BC) stem/progenitor cells, also known as tumour initiating cells, are a small, difficult to target population within the tumour responsible for metastasis and therapy resistance. MYBL2 overexpression is commonly linked to poor prognosis and metastasis but its role in BC stem/progenitor cells remains elusive. To determine the role of MYBL2 in BC stem/progenitor cells we generated TNBC cell lines with inducible MYBL2 downregulation. Our findings reveal that elevated MYBL2 is essential for self-renewal, DNA repair, and replication stress response in these cells and lowering MYBL2 impairs stemness and self-renewal both *in vitro* and *in vivo*. Accordingly, our functional and mechanistic analyses indicate that high-MYBL2 stem/progenitor cells exhibit increased sensitivity to Polθ inhibitors which is lost upon MYBL2 downregulation due to transcriptional suppression. Combining Polθ inhibitors with ATR inhibitors further enhances this sensitivity, *in vitro* and ex vivo. This study thus identifies Polθ/ATR inhibition as a synthetic lethality strategy to eradicate BC stem/progenitor cells and underscores MYBL2 expression as a biomarker for patient stratification in this treatment.

## Introduction

Breast cancer (BC) is the most diagnosed cancer worldwide and remains a leading cause of death in females [1]. The disease is extremely heterogeneous and is classified into different molecular subtypes based on expression of different cellular receptors. Triple-negative breast cancers (TNBC) lack expression of oestrogen (ER), progesterone (PR) and human epidermal growth factor receptor-2 (HER2) [2] and accounts for 10-20% of all BCs [3]. When compared to other BC subtypes, overall survival of patients is reduced [4], disease relapse is higher and metastasis often occurs within 5 years. At present, patients are commonly treated using a combination of neoadjuvant immuno-/chemotherapy followed by surgery but success rates are low in the absence of a complete pathological response and disease will reoccur due to therapy resistance. Thus, there is a need to identify new therapeutic targets to develop better treatments aimed at improving the outlook for TNBC patients [5, 6]. Several studies have provided evidence that therapy resistance in TNBC is mediated via an increased proportion of BC stem/progenitor cells when compared to other tumour subtypes [7-9].

Gene expression signatures are highly dysregulated in many solid tumours, including BC [10]. Current research focuses on identifying and studying biomarkers of metastasis, treatment resistance and disease recurrence to inform treatment decisions and increase patient survival. Upregulation of *MYBL2* gene expression has been associated with impaired therapeutic response [11, 12], poor relapse-free survival and overall survival in BC [12]. The upregulation of *MYBL2* is estimated to occur in 9-13% of BCs [13, 14] and has been demonstrated to be an early event in tumour initiation [15]. *MYBL2* is one of the 21 genes included in the Oncotype DX assay as an indicator of ER+ BC recurrence [16], indicating its importance clinically for disease pathogenesis. *MYBL2* (B-Myb) is a transcription factor with a widely recognised role in stem and somatic cells [17-24]. It performs multiple functions including promoting cell proliferation, maintaining pluripotency, facilitating cell cycle progression, and safeguarding genome stability [18-22, 24-26].

High *MYBL2* expression is observed in aggressive BC [27] and is associated with a high proliferative signature and genome instability [28], however studies of *MYBL2* function in the stem/progenitor compartment are lacking. Our previous work indicates a role for *MYBL2* in supporting DNA double strand break (DSB) repair in hematopoietic stem cells (HSCs) [21], and more recent studies using omics data showed that *MYBL2* transcriptionally regulates DNA damage response genes in lung adenocarcinoma [29]. One of these genes is POLQ, which encodes for DNA polymerase theta (Polθ), an enzyme required for theta-mediated end joining (TMEJ) [30, 31]. TMEJ is a back-up DNA DSB repair pathway known to be relied upon by homologous recombination (HR) deficient cells [32]. TNBC cells overexpress POLQ [33-35], which could indicate an increased reliance on its functions in DNA repair, however functional investigations have not been performed.

Here we used *in vitro* and *in vivo* models to investigate the role of *MYBL2* in TNBC using enriched stem/progenitor cells. We observed that *MYBL2* downregulation significantly impairs the proliferation and self-renewal of TNBC stem/progenitor cells, characterised by reduced tumour initiation and growth in a xenograft model. Downregulation of *MYBL2* in this compartment delays DNA DSB repair, which is associated with reduced TMEJ, mediated by lack of transcriptional regulation of POLQ by *MYBL2*. Exploiting the TMEJ dependency of high *MYBL2* tumours, we show that TNBC stem/progenitor cells enriched from patient-derived xenografts expressing high levels of *MYBL2* are sensitive to Polθ inhibition. This sensitivity can be further increased through combined inhibition of Polθ and ATR. Our data establish the importance of *MYBL2* in TNBC stem/progenitor cells and highlight *MYBL2* expression as a potential biomarker to stratify patients that would respond to treatment with Polθ inhibitors, effectively targeting tumour initiating cells to eradicate them and prevent disease recurrence.

## Results

### High *MYBL2* levels maintain stemness in populations enriched for TNBC stem/progenitor activity

*MYBL2* is known to be important for maintaining stem cell properties [36-38], however its specific role in TNBC stem/progenitor cells has not been determined. Similarly to previous findings in other stem cell populations [24, 39], we observed that *MYBL2* expression levels were significantly increased in stem/progenitor cells enriched from the total MDA-MB-231 cell population when compared to the bulk **(Figure 1A)**. This was the case for stem/progenitor cells purified via surface marker expression as well as functional assays, with anoikis resistant (AR) cells displaying the highest *MYBL2* expression and greatest tumour initiation capacity *in vivo* **(Figure S1)**. To determine the functional significance of this high *MYBL2* expression, we utilised a doxycycline inducible short hairpin RNA (shRNA) system to downregulate *MYBL2* levels in the TNBC lines MDA-MB-231 and CAL-51 **(Figure 1B)** and examined the consequences for the stem/progenitor population. Firstly, we measured the ability of TNBC cells to form mammospheres following *MYBL2* downregulation to study their proliferation and self-renewal capacity [40]. After addition of doxycycline, expression of the red fluorescent protein reporter could be observed within 24h and was maintained throughout the 5-day period of primary mammosphere formation **(Figure 1C)**. Downregulation of *MYBL2* had minimal effects on the number of primary mammospheres formed **(Figure 1D)**, however the size of the mammospheres was significantly reduced **(Figure 1C and 1E)** suggestive of a proliferation defect. To confirm this, we performed a BrdU incorporation assay and indeed determined that mammosphere cultures with downregulation of *MYBL2* contained a reduced percentage of proliferating BrdU+ cells **(Figure 1F**). *MYBL2* downregulation significantly decreased secondary mammosphere formation in the two TNBC cell lines examined **(Figure 1D)**, indicating an important role for *MYBL2* in self-renewal. To further evaluate primary mammosphere cultures we performed ALDEFLUOR staining to identify stem/progenitor cells and observed that downregulation of *MYBL2* significantly decreased the percentage of ALDEFLUOR+ cells **(Figure 1G)**. Furthermore, we determined that *MYBL2* downregulation in both mammosphere cultures and AR cells was associated with a significant reduction in gene expression of the pluripotency factors OCT4, SOX2 and NANOG **(Figure 1H)**. Overall, these data strongly indicate that *MYBL2* is required for maintaining TNBC stem/progenitor cells *in vitro*.

**Figure 1.**
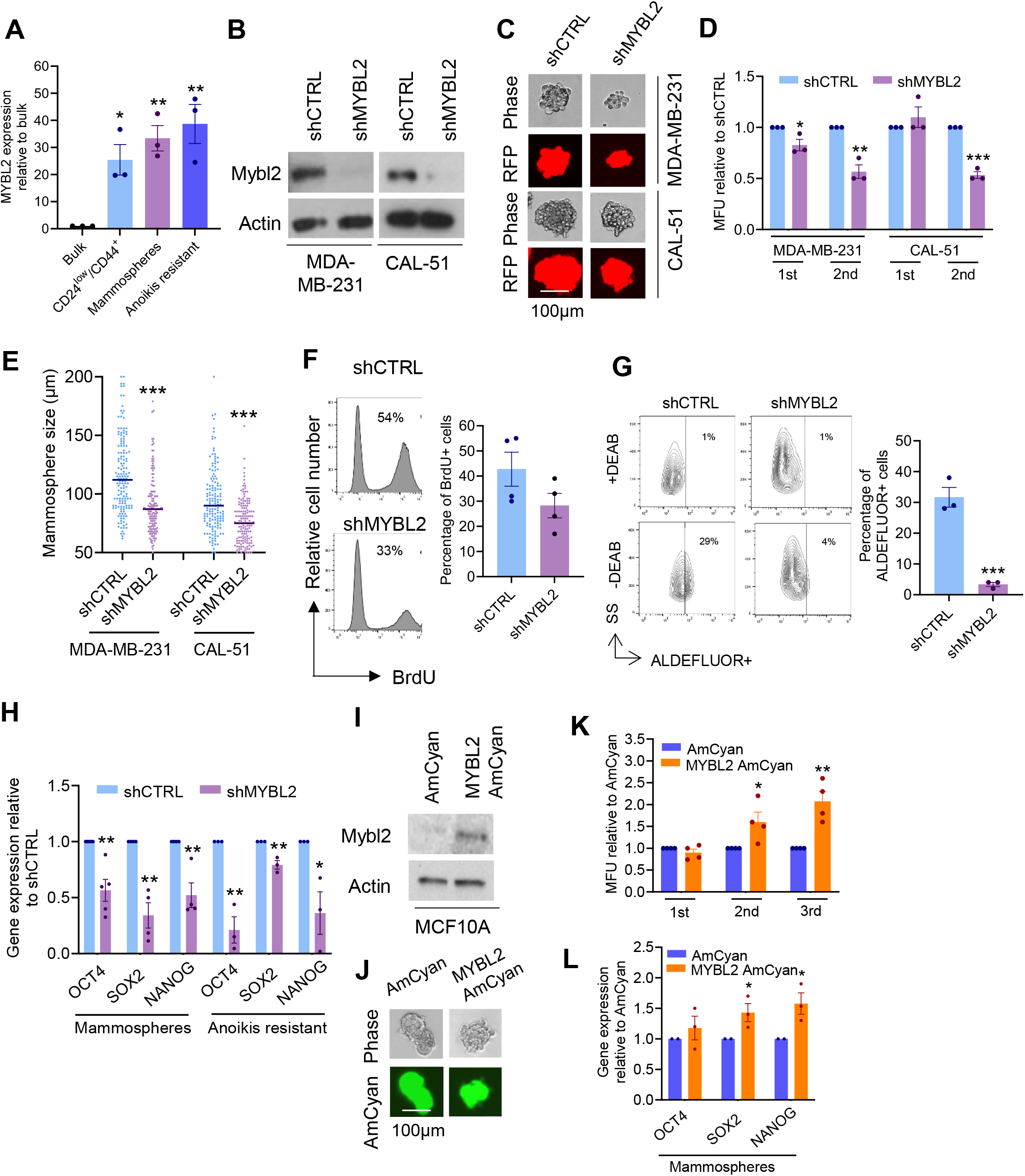
MYBL2 maintains the proliferation and self-renewal capacity of cell populations enriched for triple negative breast cancer stem/progenitor activity. (A) Breast cancer stem/progenitor cells were enriched from the bulk population of MDA-MB-231 cells using different methods and MYBL2 expression measured using qPCR. (B) MDA-MB-231 and CAL-51 cells expressing pTRIPZ Inducible Lentiviral shRNA control (shCTRL) or MYBL2 (shMYBL2) were incubated with doxycycline for 5 days and analyzed by immunoblotting. (C-E) Mammosphere formation by shCTRL and shMYBL2 MDA-MB-231 and CAL-51 cells. Representative microscopy images of primary mammospheres are shown (C), quantification of primary and secondary mammosphere numbers measured in mammosphere forming units (MFU) (D) and mammosphere size (E). (F) Primary mammosphere cultures of MDA-MB-231 cells were incubated with BrdU, stained using antibodies and analyzed by flow cytometry. (G) Primary mammosphere cultures of MDA-MB-231 cells were stained using the ALDEFLUOR assay and analyzed by flow cytometry. (H) Breast cancer stem/progenitor cells were enriched from the bulk population of shCTRL or shMYBL2 MDA-MB-231 cells as indicated and gene expression measured using qPCR. (I) MCF10A cells expressing pHAGE-2-EF1a-Linker-HA-SPA-EF1a-AmCyan (AmCyan) or pHAGE-2-EF1a-Mybl2-Linker-HA-SPA-EF1a-AmCyan (MYBL2 AmCyan) were analyzed by immunoblotting. (J-K) Mammosphere formation by MCF10A AmCyan and MYBL2 AmCyan cells. Representative microscopy images of primary mammospheres are shown (J) and quantification of mammosphere numbers measured in MFU (K). (L) Mammospheres were generated using isogenic MCF10A AmCyan and MYBL2 AmCyan cells and gene expression measured by qPCR. Plots represent data from at least three independent experiments and are shown as mean±SEM. *p ≤0.05, **p ≤0.01 and ***p ≤0.001 and obtained from a two-tailed t test.

Given that breast tumours often have multiple genetic alterations, we next wanted to investigate if *MYBL2* upregulation alone could alter the function of normal mammary stem cells to induce a BC stem cell-like phenotype. To do this, we overexpressed *MYBL2* in the non-tumourigenic mammary epithelial cell line MCF10A **(Figure 1I)** and examined the effect of this on the mammary stem cell population. The presence of cells constitutively overexpressing *MYBL2* (MYBL2 AmCyan) or the fluorescent tag alone (AmCyan) was confirmed and could be observed in mammosphere cultures **(Figure 1J)**. *MYBL2* overexpression had no effect on the number of primary mammospheres formed **(Figure 1K)** indicating that increasing *MYBL2* expression alone is not sufficient to increase the proliferation of normal mammary stem cells. In contrast, increased *MYBL2* expression did significantly increase the ability of MCF10A cells to form secondary and tertiary mammospheres **(Figure 1K)**. These data reveal that increasing *MYBL2* expression alone in mammary stem cells increases their self-renewal capacity. This is supported by gene expression analysis of primary mammosphere cultures showing that *OCT4, SOX2* and *NANOG* expression is higher when *MYBL2* is upregulated **(Figure 1L)**.

Following our *in vitro* observations, we wanted to determine if *MYBL2* was important for TNBC tumour initiation *in vivo*. To assess this, we performed a limiting dilution analysis in which NOD/Scid/IL-2Rγnull (NSG) mice received one of three different dilutions of either shCTRL or shMYBL2 bulk TNBC cells (MDA-MB-231) via subcutaneous injection **(Figure 2A)**. Animals were monitored for tumour growth over time, with the assumption being that tumours derived from cell populations containing an increased proportion of tumour initiating cells would be detectable at earlier time points. To capture the very early stages of tumour initiation, we engineered MDA-MB-231 cells to constitutively express the luciferase gene to allow *in vivo* imaging of tumour growth. Importantly, shCTRL and shMYBL2 cells exhibited similar levels of luminescence in vitro, indicating that luciferase expression was comparable between the two cell lines **(Figure S2)**. In vivo imaging revealed that TNBC cells could be detected as early as three days after injection **(Figure 2B)** and quantification of the measurements indicated shCTRL cells had increased tumour initiation capacity when compared to shMYBL2 cells **(Figure 2C)**. This was also reflected in the tumour size, most strikingly at Day 21, by which time tumours derived from 1×10^5^ shCTRL cells were significantly larger than those derived from 1×10^5^ shMYBL2 cells **(Figure 2D)**. Indeed, after 21 days animals injected with 1×10^5^ shCTRL cells had reached the license limit and tumours were excised from all animals in the cohort at this time point. MYBL2 protein downregulation was confirmed **(Figure 2E)** and the tumours physical appearance assessed **(Figure 2F)**. Tumours from animals injected with 1×10^5^ MYBL2 downregulated cells tended to be smaller in size compared to 1×10^5^ control cells **(Figure 2F+G)**. For example, the average weight of a tumour derived from 1×10^5^ shCTRL cells was 169mg whereas the average weight of a tumour derived from 1×10^5^ shMYBL2 cells was 104mg. Histological analysis of the excised tumours **(Figure 2H)** showed that 6/6 derived from shCTRL cells exhibited increased cellularity and no evidence of fibrosis, contributing to their classification as high-grade tumours. In contrast, a proportion of the tumours derived from shMYBL2 cells displayed lower cellularity (3/6) and evidence of fibrosis (2/6) resulting in classification as a lower grade tumour (3/6). These features suggest that downregulation of MYBL2 results in the formation of a less aggressive breast tumour. A limiting dilution transplant assay assessed using L-Calc (Stem Cell Technologies), estimated the stem cell frequency in shCTRL tumours was 1 in 37,585 **(Figure 2I)**, with MYBL2 downregulation decreasing this by 7-fold to 1 in 280,030. We then used the mammosphere assay to determine the functionality of stem/progenitors cells from isolated tumours ex vivo. In support of our *in vivo* data, we observed that cells from shMYBL2 tumours had reduced proliferative and self-renewal capacity when compared to cells from shCTRL tumours, as assessed by decreased primary and secondary mammosphere formation **(Figure 2J)**. In summary, we have shown that high levels of MYBL2 promote the proliferative and self-renewal activities of TNBC stem/progenitor cells.

**Figure 2.**
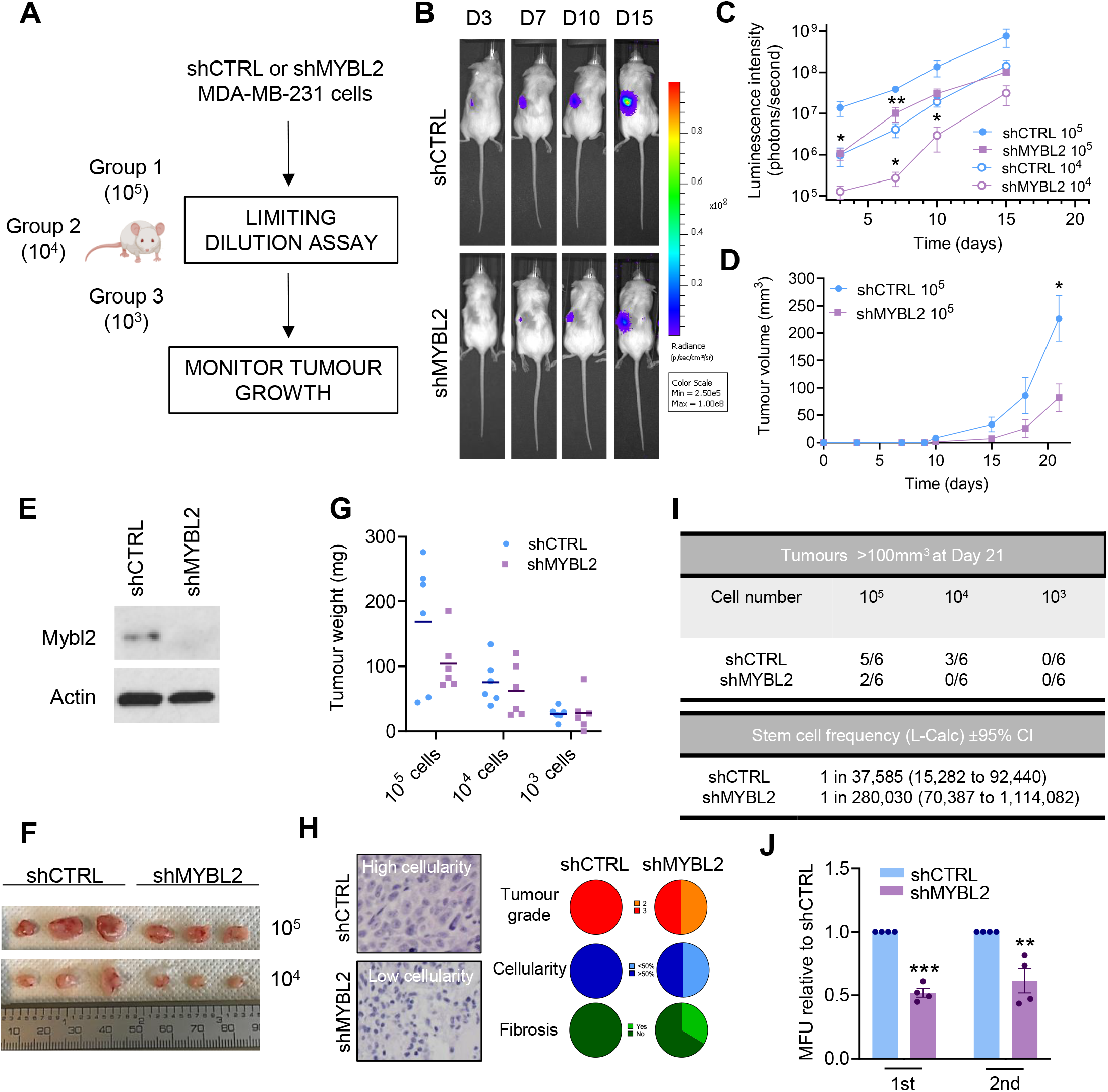
MYBL2 promotes triple negative breast tumour initiation and growth *in vivo*. (A) Schematic of the limiting dilution assay. Different numbers of shCTRL or shMYBL2 MDA-MB-231 cells were injected into the flank of NSG mice. (B) Representative IVIS images over time of animals injected with 10^4^ shCTRL or shMYBL2 MDA-MB-231 cells. (C) Quantification of IVIS images of animals injected with shCTRL or shMYBL2 MDA-MB-231 cells over the course of the experiment. (D) Tumour volume calculated from caliper measurements of tumours formed by shCTRL or shMYBL2 MDA-MB-231 isogenic cells. (E) MYBL2 protein expression in cells harvested from shCTRL or shMYBL2 tumours. (F) Photographic image of representative tumours formed by shCTRL and shMYBL2 MDA-MB-231 cells at Day 21. (G) Weight of tumours formed by shCTRL and shMYBL2 MDA-MB-231 cells at Day 21. (H) Representative images of H+E stained tumour sections and pie charts indicating the classification of tumours in terms of grade, cellularity and fibrosis. (I) Table indicating the total number of tumours detected in each experimental group and calculation of the stem cell frequency in shCTRL and shMYBL2 tumours. (J) Mammosphere formation by shCTRL and shMYBL2 MDA-MB-231 cells harvested from tumours. Quantification of mammosphere numbers is measured in mammosphere forming units (MFU). Plots represent data from six independent animals and are shown as mean±SEM. Mammosphere formation was measured using four independent tumours from each group. *p ≤0.05, **p ≤0.01 and ***p ≤0.001 and obtained from a two-tailed t test.

### *MYBL2* supports DNA double strand break repair in populations enriched for stem/progenitor cells from TNBC

We next sought to determine how high *MYBL2* expression could be maintaining the stem/progenitor cell population in TNBC. Previous work has shown that *MYBL2* has an important role in maintaining genome stability and supporting DNA DSB repair [18-22, 24], two processes which are particularly important in stem cells. Stem cells have been reported to be less susceptible to DNA damage due to increased DNA repair capacity [41, 42], perhaps due to the activation of additional pathways, which contributes to their resistance to DNA damaging chemotherapeutics. To compare the DSB repair capacity of the bulk and stem/progenitor populations after *MYBL2* downregulation, we monitored the kinetics of irradiation-induced foci (IRIF) of 53BP1, a well characterised marker of DSBs [43]. The bulk cell population was much slower at repairing irradiation-induced DSBs when compared to the stem/progenitor population **(Figure 3A-D**). This was particularly evident 5h post-IR when stem/progenitor cells showed 53BP1 clearance, whereas the bulk cell population did not. *MYBL2* downregulation slowed 53BP1 clearance in both the bulk and stem/progenitor populations and resulted in the persistence of 53BP1 foci at time points as late as 24-72h post-IR **(Figure 3C+D and S3A+B**) and elevated IR-induced genome instability **(Figure 3E and S3C)** in the stem/progenitor population. To gain further insight into this phenotype, we measured IRIF of proteins involved in the two main DSB repair pathways: non-homologous end joining (NHEJ) and HR. We observed a modest increase in IRIF of the NHEJ protein RIF1 5h after treatment in *MYBL2* downregulated stem/progenitor cells derived from MDA-MB-231 cells but no change in CAL-51 derived stem/progenitor cells **(Figure 3F and S3D)**. The most striking phenotypes were a reduction in IRIF of the HR proteins BRCA1 and RAD51 in S/G2 cells **(Figure 3G+H and S3E+F)**. This was attributed to a decrease in the overall protein levels of BRCA1 and RAD51 in stem/progenitor cells with downregulation of *MYBL2* **(Figure 3I)**. In contrast, levels of 53BP1 and RIF1 protein remained unaffected following *MYBL2* downregulation **(Figure 3I)**, indicating that high levels of *MYBL2* promote the expression and recruitment of proteins involved in HR.

**Figure 3.**
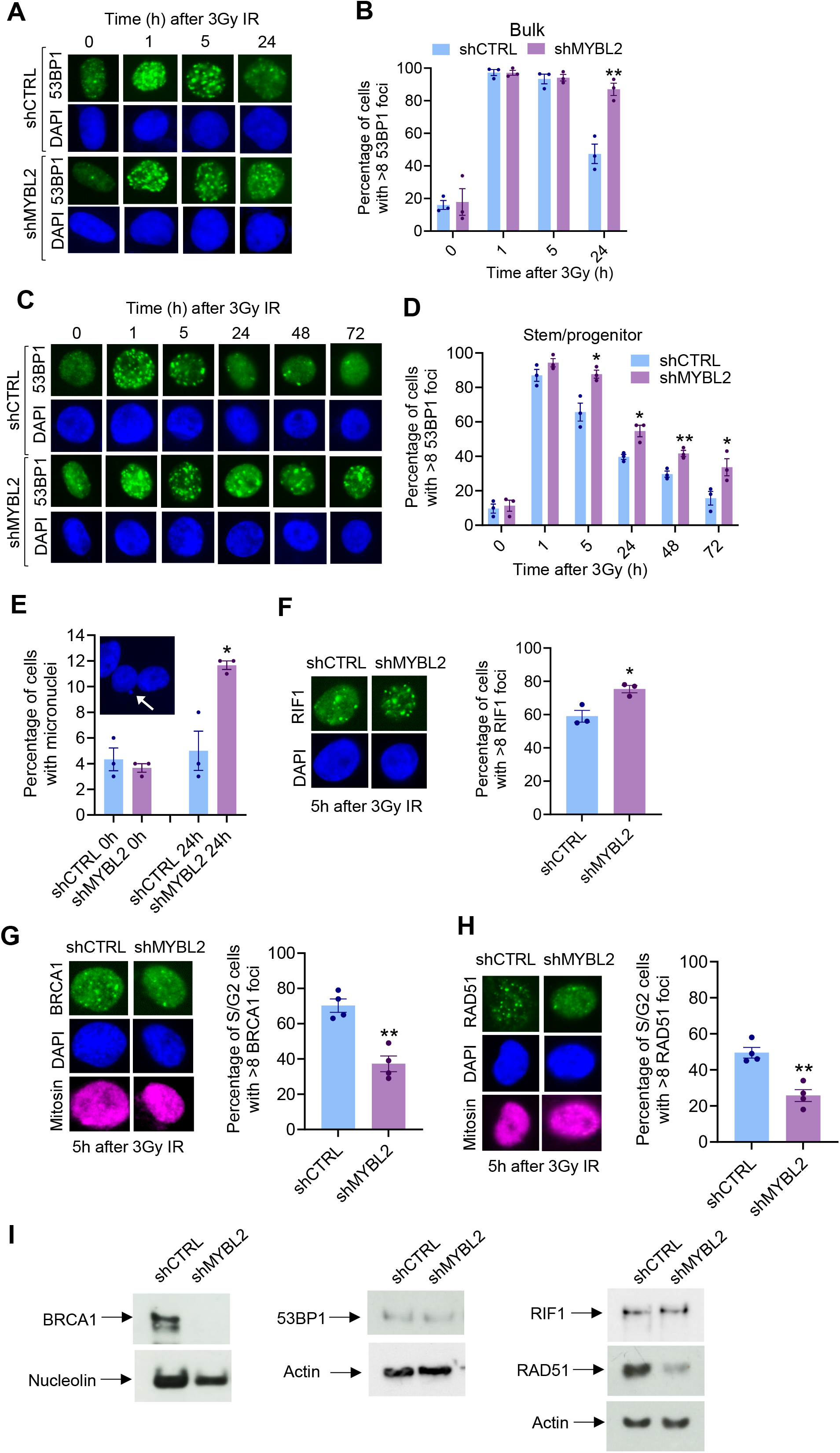
MYBL2 supports DNA double strand break repair in cell populations enriched for triple negative breast cancer stem/progenitor cell activity derived from bulk MDA-MB-231 cells. (A-D) Bulk MDA-MB-231 cells (A+B) or mammosphere cultures (C+D) expressing pTRIPZ Inducible Lentiviral shRNA control (shCTRL) or MYBL2 (shMYBL2) were incubated with doxycycline for 5 days, exposed to ionizing radiation (IR) and immunostained with antibodies to 53BP1 at the indicated time points. Representative microscopy images are shown (A+C) and quantification of 53BP1+ cells (B+D). (E) Mammosphere cultures were treated with doxycycline as in (C+D), exposed to IR and micronuclei formation assessed 24h later. (F-H) Mammosphere cultures were treated with doxycycline as in (C+D), exposed to IR and immunostained with antibodies to RIF1 (F), BRCA1 and mitosin (G) or RAD51 and mitosin (H). For each panel representative microscopy images and foci quantification are shown. (I) Mammosphere cultures were treated with doxycycline as in (C+D) and indicated proteins analyzed by immunoblotting. Plots represent data from at least three independent experiments and are shown as mean±SEM. *p ≤0.05, **p ≤0.01 and ***p ≤0.001 and obtained from a two-tailed t test.

### *MYBL2* promotes the activity of Polθ in stem/progenitor enriched populations from TNBC

Considering that the stem/progenitor population display faster 53BP1 clearance when compared to the bulk cell population, we were prompted to investigate whether stem/progenitor cells use alternative DNA repair pathways to maintain this phenotype and whether these were affected under conditions of *MYBL2* downregulation. One of the main back-up HR pathways is single-strand annealing (SSA), an error prone repair process involving RAD52 [44, 45]. Inhibition of RAD52 is known to reduce the proliferation of *BRCA* deficient cells [46, 47], and therefore represents an attractive target for the treatment of *BRCA* deficient BCs. The other main back up HR pathway is TMEJ, another error-prone repair process involving DNA polymerase theta (Polθ) [48]. Similarly to RAD52, Polθ inhibition is also under investigation as a treatment for HR-deficient cancers [49, 50]. To determine the potential effects of *MYBL2* loss on SSA and TMEJ in the stem/progenitor compartment, we measured IRIF of RAD52 and Polθ. We observed an increase in IRIF of RAD52 5h after treatment in *MYBL2* downregulated cells **(Figure 4A and S4A)**, perhaps indicating an increased reliance on the SSA pathway to repair DNA damage. In contrast, there was a reduction in IRIF of Polθ in S/G2 cells following *MYBL2* downregulation **(Figure 4B + S4B)**, suggesting that similarly to the classical HR proteins, recruitment of proteins involved in TMEJ is also reduced when *MYBL2* is downregulated.

**Figure 4.**
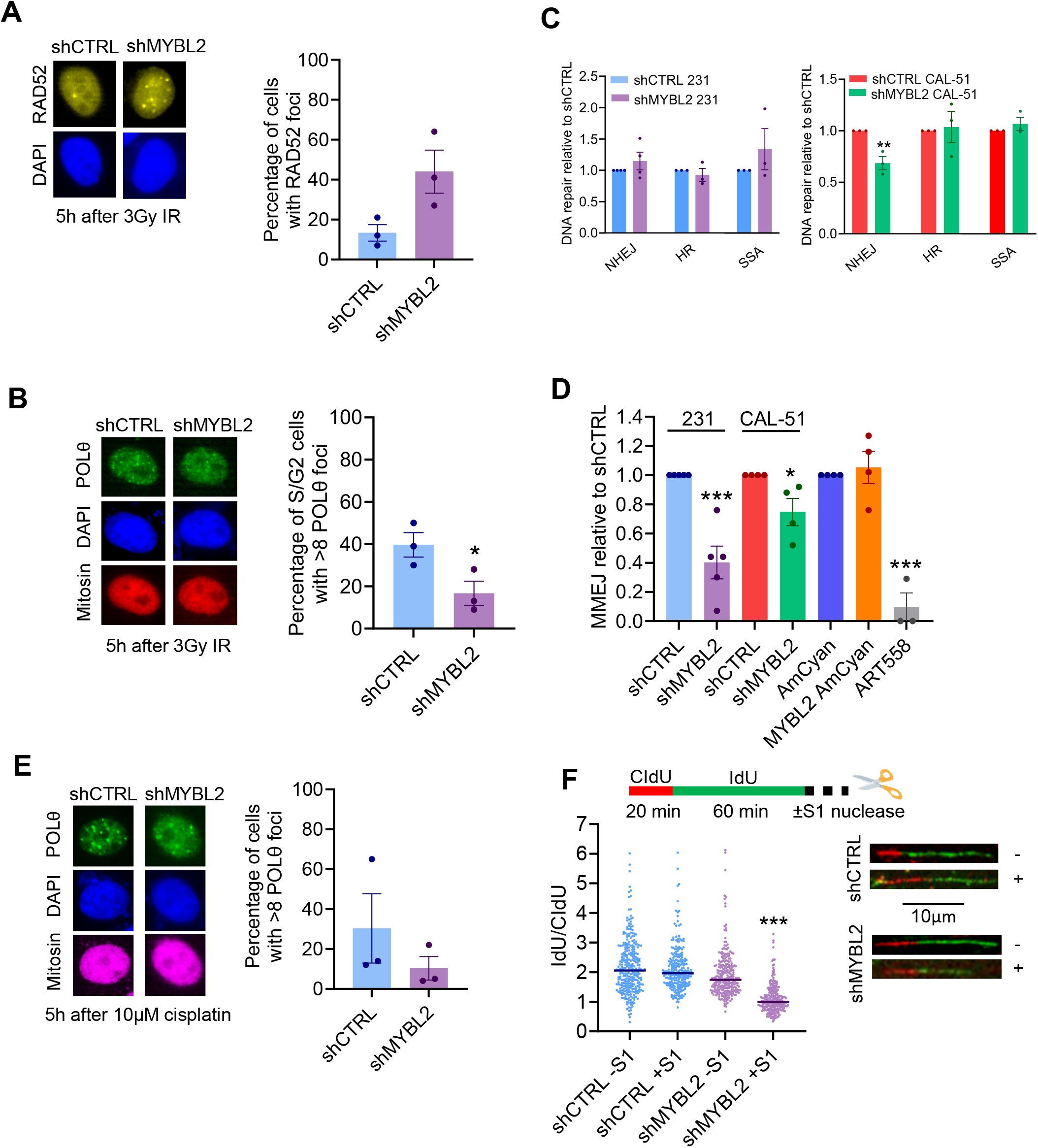
High levels of MYBL2 promotes the activity of Polθ in cell populations enriched for triple negative breast cancer stem/progenitor cell activity. (A+B) Anoikis resistant cells were isolated from bulk MDA-MB-231 cultures expressing pTRIPZ Inducible Lentiviral shRNA control (shCTRL) or MYBL2 (shMYBL2). Cells were transfected with RAD52-YFP (A) or Polθ-FLAG (B) plasmids, exposed to ionizing radiation (IR) and imunostained with antibodies to mitosin (A) or FLAG and mitosin (B). For each panel representative microscopy images and foci quantification are shown. (C) Anoikis resistant MDA-MB-231 and CAL-51 shCTRL and shMYBL2 cells were transfected with NHEJ, HR or SSA NanoLuciferase reporter plasmids. Cells performing DNA repair via each specific pathway were quantified via measurement of luminescence. (D) Anoikis resistant shCTRL and shMYBL2 cells isolated from bulk MDA-MB-231 or CAL-51 cultures or MCF10A cells expressing pHAGE-2-EF1a-Linker-HA-SPA-EF1a-AmCyan (AmCyan) or pHAGE-2-EF1a-Mybl2-Linker-HA-SPA-EF1a-AmCyan (MYBL2 AmCyan) were transfected with an TMEJ NanoLuciferase report plasmid. Cells performing DNA repair via TMEJ were quantified via measurement of luminescence. (E) Anoikis resistant shCTRL and shMYBL2 cells isolated from bulk MDA-MB-231 cultures were transfected with Polθ-FLAG plasmid, treated with cisplatin and immunostained with antibodies to FLAG and mitosin. Representative microscopy images and foci quantification are shown. (F) Anoikis resistant shCTRL and shMYBL2 cells isolated from bulk MDA-MB-231 cultures were pulsed with CldU and IdU and treated with S1 nuclease. DNA was visualized by immunostaining with antibodies to CIdU and IdU and IdU:CIdU ratio was quantified in two independent experiments. ***p ≤0.001 and obtained from a Mann-Whitney test. All other plots represent data from at least three independent experiments and are shown as mean±SEM. *p ≤0.05, **p ≤0.01 and ***p ≤0.001 and obtained from a two-tailed t test.

To ascertain the functional significance of our IRIF measurements, we used luminescence-based DNA repair reporter assays [49, 51] to measure the efficiency of HR, NHEJ, SSA and TMEJ DNA repair pathways. These assays revealed that despite changes in protein levels and/or recruitment, DNA repair by HR, NHEJ and SSA in stem/progenitor enriched population from the MDA-MB-231 cell line was unaffected by *MYBL2* downregulation **(Figure 4C)**. Similar results were observed in the CAL-51 cell line for HR and SSA, however a decrease in NHEJ could also be observed **(Figure 4C)**. Overall, these findings indicate that the amount of BRCA1/RAD51 present in the cell is sufficient to repair the exogenous damage using HR. The HR proficiency of *MYBL2* downregulated stem/progenitor cells was also reflected in their insensitivity to the PARP inhibitor olaparib **(Figure S4C)**. However, loss of *MYBL2* did impair the ability of stem/progenitor cells to repair DNA by TMEJ **(Figure 4D)**, indicating that use of this pathway could be influenced by *MYBL2* levels. We also observed that levels of TMEJ were very low in mammary stem cells from the non-tumourigenic cell line MCF10A when compared to the TNBC lines and was unaffected by *MYBL2* overexpression **(Figure 4D and S4D)**.

To further examine the relationship between *MYBL2* and Polθ, we wanted to determine if our findings were specific to IR-induced DNA damage and if other Polθ-mediated functions were affected under conditions of *MYBL2* downregulation in stem/progenitor cells. To this end, *MYBL2* dowregulated stem/progenitor cells treated with cisplatin also displayed reduced Polθ recruitment to damage sites **(Figure 4E and S4E)**, suggesting our findings could also be applicable to other DNA damaging agents. Aside from TMEJ, Polθ has also been shown to have an important role in processing single-stranded DNA (ssDNA) gaps in *BRCA* deficient cells [52, 53]. We have shown that *MYBL2* downregulation reduces BRCA1 protein levels in stem/progenitor cells **(Figure 3I)** without affecting DNA repair by HR **(Figure 4C)** but had not determined if other BRCA-dependent functions were affected, for example the emergence of ssDNA gaps. To study this, we labelled replication forks with nucleotide analogues and digested them using the S1 nuclease to determine if replication tract length was shortened **(Figure 4F)**. Replication tracts in stem/progenitor cells with lower levels of *MYBL2* showed significant shortening in the presence of S1 nuclease, whereas those with high *MYBL2* levels were resistant to the S1 nuclease treatment **(Figure 4F)**. This demonstrates that post-replicative ssDNA gaps are present in *MYBL2* deficient stem/progenitor cells in the presence of normal HR, indicating that the reduction in BRCA1 protein observed under conditions of *MYBL2* downregulation affects the resolution of ssDNA gaps which would usually be sealed by Polθ.

### *MYBL2* drives expression of genes that promote homologous recombination and theta-mediated end joining

*MYBL2* is a well characterised transcription factor known to regulate genes important for cell cycle control [54-56], survival [57] and differentiation [38]. However, its effects on gene expression in BC stem/progenitor cells remains unexplored, prompting us to hypothesise that *MYBL2* could be regulating the transcription of genes important for the DNA damage response such as BRCA1, RAD51 and POLQ (Polθ). To investigate this, we performed qPCR analysis of gene expression levels in stem/progenitor cells isolated from MDA-MB-231 cell cultures in which levels of *MYBL2* had been downregulated. We observed that *MYBL2* downregulation had no effect on the level of 53BP1 and RIF1 gene expression in stem/progenitor cells **(Figure 5A)**. In contrast, expression levels of BRCA1, RAD51 and POLQ were significantly reduced following *MYBL2* downregulation, which in the case of POLQ was independent of the method used to isolate the stem/progenitor cells **(Figure 5A+B)**. Overexpression of *MYBL2* in the normal mammary cell line MCF10A had no effect on the expression levels of 53BP1, RIF1, BRCA1, RAD51 and POLQ **(Figure S5A)**. To further examine the correlation between *MYBL2* levels and expression of BRCA1, RAD51 and POLQ in BC we analysed omics data available from The Cancer Genome Atlas (TCGA) via cBioPortal [58-60]. TCGA samples with RNA sequencing data (Firehose Legacy, n=1108) were classified as *MYBL2*high or *MYBL2*low based on *MYBL2* mRNA expression. Using this stratification criteria, we observed that *MYBL2*high BCs were associated with high expression levels of *BRCA1, RAD51* and *POLQ* **(Figure 5C)**. Through analysis of data available through the Cancer Cell Line Encyclopedia (CCLE) we have identified that this association is also present in BC cell lines, demonstrating the consistency of our findings in both cell lines as well as primary BC samples **(Figure S5B)**. Next, we investigated whether *MYBL2* was directly regulating the expression of *BRCA1, RAD51* and *POLQ*. To study this, we performed chromatin immunoprecipitation (ChIP) in the bulk population of MDA-MB-231 cells to determine enrichment of MYBL2 at genomic regions previously identified using ChIP-seq in HepG2 cells [61]. We observed a strong enrichment of MYBL2 at regions in the *CCNB1* gene **(Figure 5D)**, which is a known target of MYBL2 [55, 62, 63]. Furthermore, we could also see some enrichment of MYBL2 at regions in the POLQ, BRCA1 and RAD51 genes in the bulk cell population **(Figure 5D)**, as well as in the POLQ gene in AR-cells **(Figure 5E)**. To further confirm that MYBL2 was transcriptionally regulating POLQ, we generated constructs containing MYBL2 mutant forms with either the N-terminal DNA binding domain or the C-terminal domain truncated. These constructs were transfected into the shMYBL2 MDA-MB-231 cell lines that were doxycycline treated to eliminate the endogenous MYBL2. These data reveal that the N-terminal domain of MYBL2 is required for POLQ expression **(Figure 5F)** as well as functional TMEJ **(Figure 5G)**. Overall, these data establish that MYBL2 has a role in the transcriptional activation of specific genes associated with the DNA damage response.

**Figure 5.**
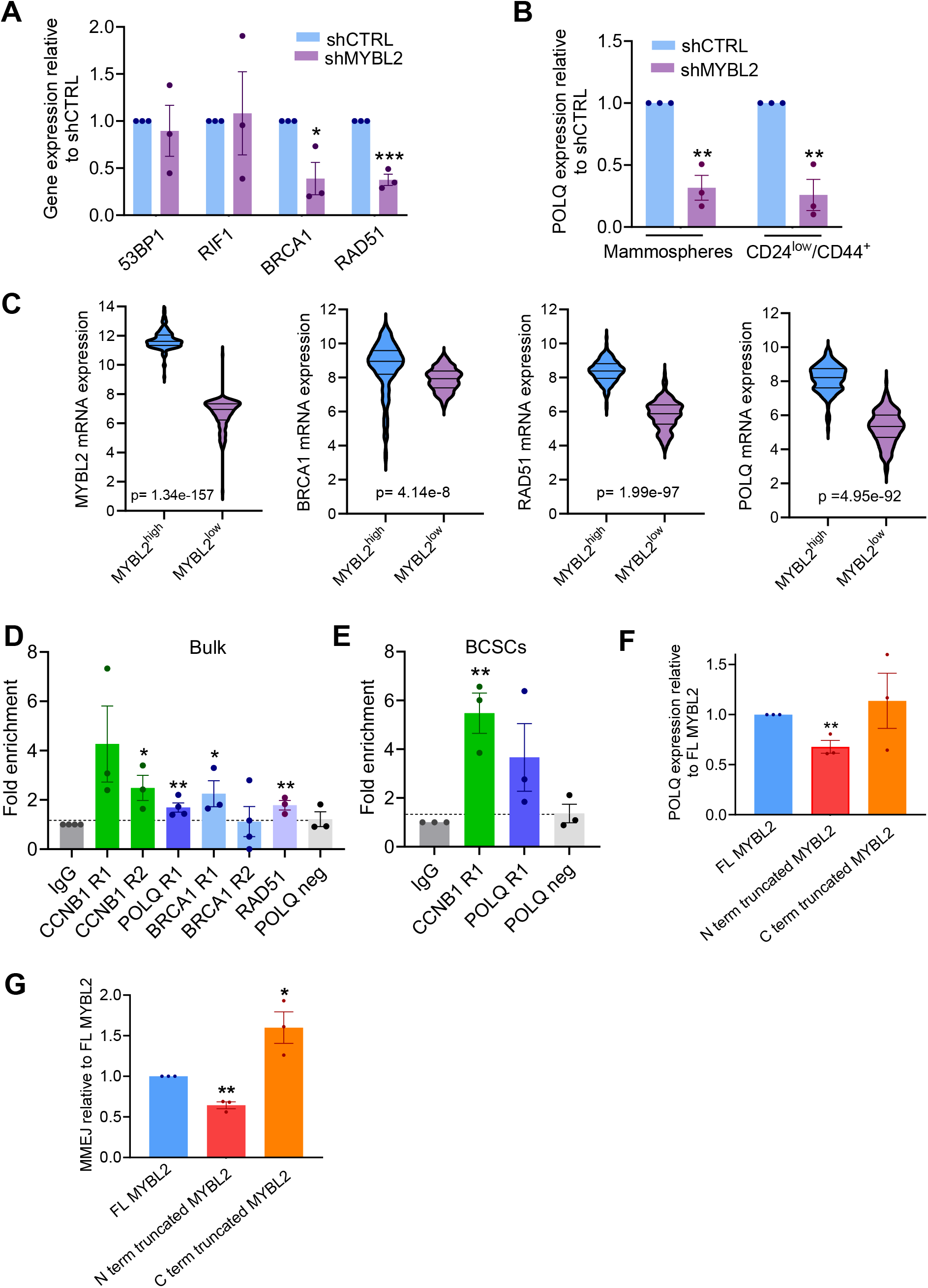
MYBL2 drives expression of genes that promote homologous recombination and theta-mediated end joining in cell populations enriched for triple negative breast cancer stem/progenitor cell activity. (A) Breast cancer stem cells were enriched from the bulk population of shCTRL or shMYBL2 MDA-MB-231 cells via mammosphere formation (A+B) or CD24/44 expression (B) and gene expression measured using qPCR. (C) mRNA expression of DNA repair genes in breast cancer patients separated into two groups by MYBL2 mRNA expression. (D+E) Chromatin was isolated from bulk (D) or anoikis resistant (E) MDA-MB-231 cells and MYBL2 ChIP was performed. Data show the enrichment of the indicated genes relative to the input and IgG control. (F-G) MDA-MB-231 cells expressing pTRIPZ Inducible Lentiviral shRNA MYBL2 (shMYBL2) and pHAGE-2-EF1a-Mybl2-L-HA-SPA-EF1a-AmCyan (FL MYBL2), or pHAGE-2-EF1a-N term truncated Mybl2-L-HA-SPA-EF1a-AmCyan (N term truncated MYBL2) or pHAGE-2-EF1a-C term truncated Mybl2-L-HA-SPA-EF1a-AmCyan (C term truncated MYBL2) were incubated with doxycycline and POLQ expression measured by qPCR (F) or transfected with an TMEJ NanoLuciferase DNA-repair reporter plasmid to measure TMEJ (G). All plots represent data from at least three independent experiments and are shown as mean±SEM. *p ≤0.05, **p ≤0.01 and ***p ≤0.001 and obtained from a two-tailed t test.

### TNBC stem/progenitor cells with high levels of *MYBL2* can be targeted using a Polθ inhibitor

We have found that MYBL2 influences the expression and/or recruitment of BRCA1, RAD51 and Polθ, proteins important for distinct DSB repair pathways (HR and TMEJ). Our functional studies show that under conditions of MYBL2 downregulation stem/progenitor cells are still able to maintain efficient HR, but not TMEJ **(Figure 4C+D)**. As there are no specific MYBL2 inhibitors available, our focus turned to Polθ and the TMEJ pathway as a potential way to target stem/progenitor cells and induce cell death. Our hypothesis being that stem/progenitor cells with high MYBL2 levels (shCTRL) would be susceptible to Polθ inhibition, and that this sensitivity would be lost in MYBL2 downregulated stem/progenitor cells (sh*MYBL2*). To investigate this, we isolated AR cells from the bulk population of TNBC cell lines (MDA-MB-231 and CAL51) expressing doxycycline inducible control and MYBL2 shRNA and monitored their survival in the presence of the Polθ inhibitor ART558 [49]. After 10 days, we observed that micromolar concentrations of ART558 were able to induce cell death in shCTRL cells, whereas a significantly higher percentage of shMYBL2 cells remained alive following the treatment **(Figure 6A)**. To further evaluate the effects of Polθ inhibition on stem/progenitor cell viability, we performed mammosphere assays in the presence of ART558. ART558 treatment significantly reduced the number of mammospheres formed by shCTRL cells when compared to the DMSO control **(Figure 6B)**. In contrast, ART558 had no effect on mammosphere forming ability of shMYBL2 cells in the two TNBC cell lines tested **(Figure 6B)**. Importantly, inhibition of Polθ in non-tumourigenic mammary stem cells had little or no effect on the ability of AR cells to form colonies **(Figure 6C)** or primary and secondary mammosphere formation **(Figure 6D)**. This implies Polθ inhibition could be used to specifically target triple negative stem/progenitor cells without affecting the proliferation and self-renewal of normal mammary stem cells.

**Figure 6.**
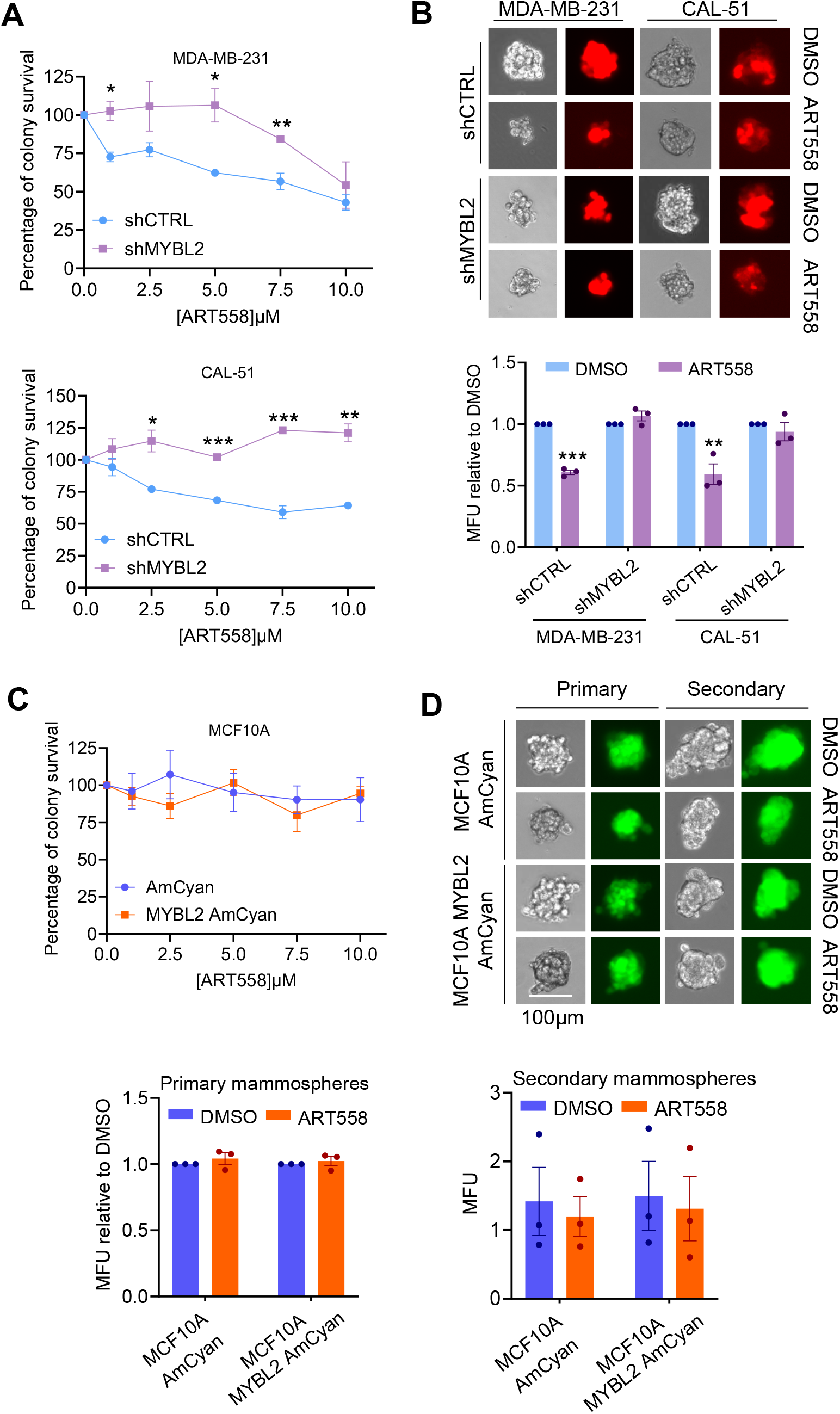
Cell populations enriched for triple negative breast cancer stem/progenitor cell activity with high levels of MYBL2 can be targeted with a Polθ inhibitor. (A) Anoikis resistant cells were isolated from bulk MDA-MB-231 or CAL-51 cultures expressing pTRIPZ Inducible Lentiviral shRNA control (shCTRL) or MYBL2 (shMYBL2). Cells were treated with ART558 and doxycycline and left for 10 days to form colonies. Colonies were stained with methylene blue and counted. (B) Mammosphere formation by shCTRL and shMYBL2 MDA-MB-231 and CAL-51 cells in the presence of 2.5µM ART558. Representative microscopy images of primary mammospheres are shown and quantification of mammosphere numbers measured in mammosphere forming units (MFU). (C) Colony formation by anoikis resistant MCF10A cells expressing pHAGE-2-EF1a-L-HA-SPA-EF1a-AmCyan (AmCyan) or pHAGE-2-EF1a-Mybl2-L-HA-SPA-EF1a-AmCyan (MYBL2 AmCyan) in the presence of ART558. Colony formation was determined as in (A). (D) Mammosphere formation by MCF10A AmCyan and MYBL2 AmCyan cells in the presence of 2.5µM ART558. Representative microscopy images and quantification of mammosphere numbers measured in MFU as shown. All plots represent data from at least three independent experiments and are shown as mean±SEM. *p ≤0.05, **p ≤0.01 and ***p ≤0.001 and obtained from a two-tailed t test.

The reduction in stem/progenitor cell survival observed in shCTRL cells following ART558 treatment prompted us to investigate whether combining this therapy with other DNA damaging agents could increase its efficacy. When ART558 was combined with PARP inhibition, Wee1 inhibition or ionising radiation there was no significant change in stem/progenitor cell survival when measured by mammosphere formation **(Figure 7A)**. However, combining ART558 with AZ20, an inhibitor of ATR kinase, did increase its efficacy **(Figure 7A)** and sensitivity was again lost in shMYBL2 stem/progenitor cells **(Figure 7B)**. The effect in shCTRL stem/progenitor cells was reproducible in cells isolated from tumours *ex vivo* **(Figure 7C)** and when alternative clinical grade inhibitors of Polθ (ART6043) and ATR (ART0380) were used **(Figure 7D)**. To determine the significance of Polθ inhibition as a therapeutic strategy for TNBCs with high levels of MYBL2, we utilised cells from a patient derived xenograft (PDX) expressing high levels of MYBL2. Primary mammosphere formation by the PDX expressing high levels of MYBL2 was decreased with ART558 treatment when compared to the DMSO control, confirming that Polθ inhibition can reduce the proliferation of primary-derived TNBC stem/progenitor cells **(Figure 7E)**. Primary mammosphere formation by cells from the PDX model was further decreased following combined treatment with ART558 and AZ20 **(Figure 7E)**, indicating that dual inhibition of Polθ and ATR kinase is an effective way to inhibit stem/progenitor cells proliferation in TNBCs with high levels of MYBL2. Overall, our data support the notion that Polθ inhibition in combination with increased replication stress will be an effective way to therapeutically target the stem/progenitor cell compartment in TNBC.

**Figure 7.**
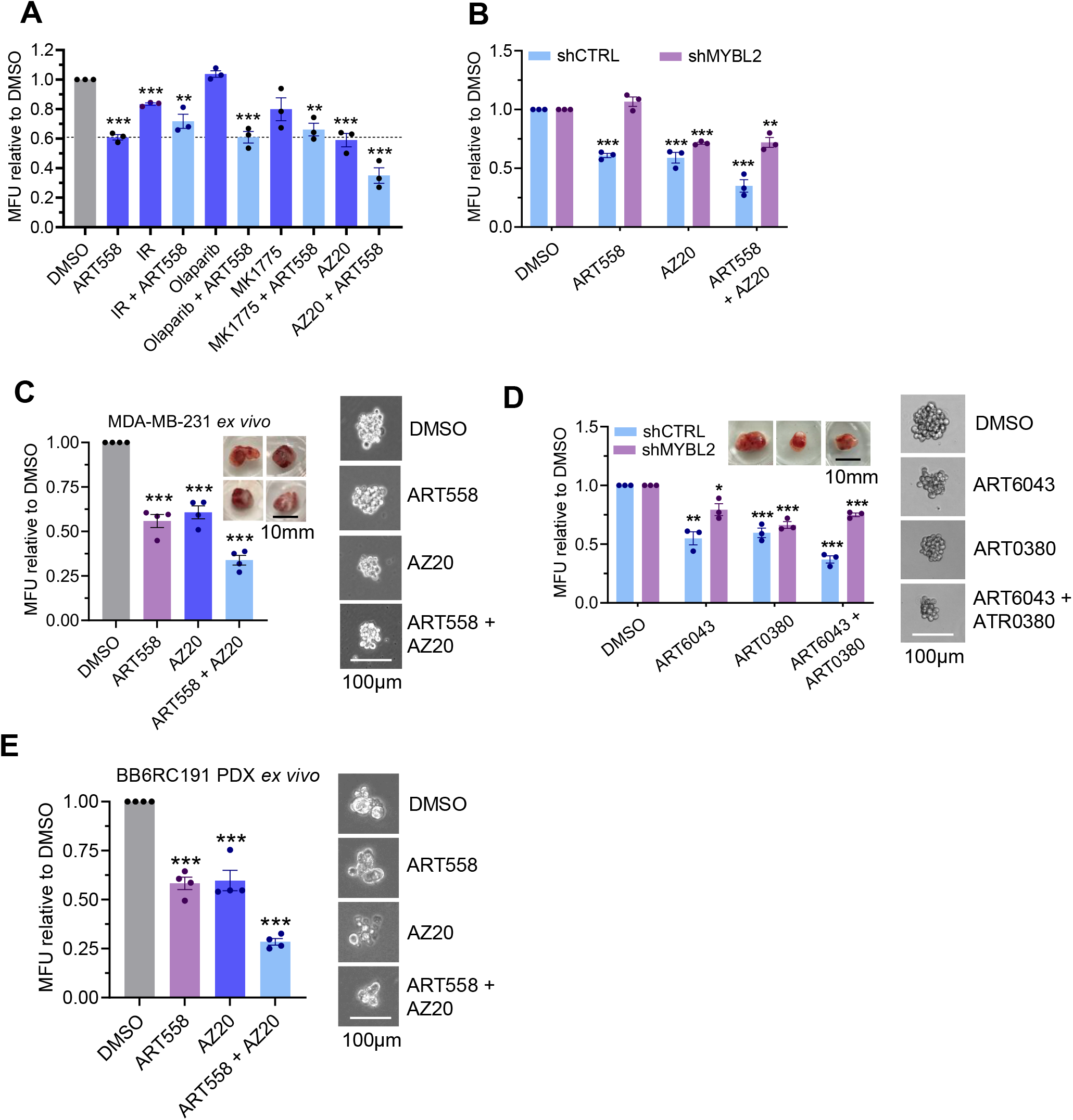
Combining Polθ and ATR inhibition enhances the killing of cell populations enriched for triple negative breast cancer stem/progenitor cell activity with high levels of MYBL2. (A) Mammosphere formation by shCTRL MDA-MB-231 cells in the presence of 2.5µM ART558 alone or in combination with different DNA damaging agents or replication stress inducers (3Gy IR, 1µM Olaparib, 50nM MK1775, 100nM AZ20). Quantification of mammosphere numbers measured in mammosphere forming units (MFU) is shown. (B-D) Mammosphere formation by shCTRL or shMYBL2 MDA-MB-231 cells in the presence of Polθ and/or ATR inhibitors. 2.5µM ART558 and/or 100nM AZ20 *in vitro* (B) and *ex vivo* (C). 2.5µM ART6043 and/or 2.5µM ART0380 clinical grade inhibitors *ex vivo* (D). Quantification of mammosphere numbers measured in mammosphere forming units (MFU) is shown, as well as photographic images of the tumours and representative microscopy images of primary mammospheres. (E) *Ex vivo* mammosphere formation by cells isolated from a patient derived xenograft in the presence of 2.5µM ART558 and/or 100nM AZ20. Representative microscopy images and quantification of mammosphere numbers measured in MFU as shown. All plots represent data from at least three independent experiments and are shown as mean±SEM. *p ≤0.05, **p ≤0.01 and ***p ≤0.001 and obtained from a two-tailed t test.

## Discussion

BC is the most diagnosed cancer among women [64, 65], with TNBC being the most aggressive subtype. This is reflected in the lower overall survival of TNBC patients when compared to the more indolent luminal subtypes [4, 66]. 90% of deaths in BC patients result from therapy resistance and disease metastasis [67], two properties associated with cancer stem/progenitor cells (also known as tumour initiating cells) [68]. Despite the importance of cancer stem/progenitor cells to BC pathogenesis, there is relatively little known about their precise origin and how they are maintained alongside the bulk cell population in breast tumours. Due to this lack of knowledge, there are currently no treatments specifically targeting tumour initiating cells. To facilitate this, more studies into the mechanisms supporting this population are required. This would allow the identification of cellular vulnerabilities to enable patient stratification and the development of new treatments to ultimately increase therapy effectiveness.

Changes in gene expression are critical to cancer initiation and progression, and as such cancer cells have been reported to have a distinct molecular signature [10]. Amplification of the *MYBL2* gene is a feature of many different solid tumours, including BC [17]. The TNBC subtype remains the most difficult to treat and is often associated with the highest *MYBL2* expression [27]. Physiologically, *MYBL2* also plays an important role in stem cells [36-38], however its role in cancer stem cells is less well defined. In this study, we uncover a role for *MYBL2* in maintaining essential properties of TNBC stem/progenitor cells including stemness and elevated DNA repair capacity. To this end, the ability of stem/progenitor cells to repair DNA damage and maintain their stem cell phenotype are thought to be key properties allowing them to survive and contribute to chemoresistance and disease recurrence in patients [69-72]. More specifically, we have found that TNBC stem/progenitor cells expressing high levels of *MYBL2* are sensitive to inhibition of Polθ, a protein important for DNA DSB repair by TMEJ [48].

Similarly to *MYBL2*, upregulation of POLQ is also associated with poor clinical outcome in BC and is most common in TNBCs [33, 34]. It represents a promising cancer treatment target due to its low expression levels in healthy mammary cells relative to BC cells. In support of this, the TNBC cell line MDA-MB-231 used in this study has been shown to express three times more POLQ at the mRNA level when compared to the untransformed mammary epithelial cell line MCF10A [73]. Previous work highlights the importance of Polθ to MDA-MB-231 cell function as siRNA-mediated knockdown reduced cell proliferation, viability and invasion capacity [74]. In addition, it has been observed that some BC cell lines express as much as eighteen times more POLQ mRNA when compared to normal mammary cells [73], implying that some TNBCs may be even more sensitive to Polθ inhibition than those used in this work.

Many studies to date have focussed on the utility of Polθ inhibitors in the context of BRCA deficiency [49, 52, 53], under the premise that cancer cells with insufficient HR use Polθ dependent TMEJ as a compensatory mechanism to retain genome stability. In contrast to this, we have found that stem/progenitor cells can use TMEJ in the context of HR proficiency. Additionally, we have found that in the absence of *MYBL2* both HR and TMEJ genes are downregulated. Although BRCA1 and RAD51 protein levels and recruitment to damage sites are reduced in stem/progenitor cells with lower levels of *MYBL2*, they remain HR proficient, PARP inhibitor insensitive and only show a decrease in functional DNA repair by TMEJ. To our knowledge these observations represent a unique finding of induced *BRCA* deficiency without effects on classical HR. However, *MYBL2* dowregulated stem/progenitor cells do have increased post-replicative ssDNA gaps, which are known to be filled by Polθ in the absence of *BRCA1/2* [53], suggesting that some BRCA dependent functions are impaired.

More recently, the use of Polθ inhibitors has been expanded to HR proficient colorectal, colon, lung and bladder cancers where it was shown to be effective when combined with radiation therapy [75, 76]. Additionally, loss of POLQ in colon cancer cells increases their sensitivity to replication stress inducing agents [35], suggesting that high POLQ expression is advantageous to tumour cells to provide a proliferative advantage. Similarly to these studies in other cancer types, we could observe a moderate but reproducible sensitivity to Polθ inhibition alone in stem/progenitor cells with high levels of *MYBL2*, which is enhanced when combined with an ATR kinase inhibitor to induce replication stress. An important further direction would be to determine if Polθ inhibition in combination with other replication stress inducers or DNA damaging agents could be an effective way to treat TNBCs with high levels of *MYBL2*.

We have shown that *MYBL2* regulates DNA repair gene expression in bulk and stem/progenitor cells at the level of transcription. Binding of MYBL2 to the promotor regions of *BRCA1, RAD5*1, and *POLQ* appears to be required to maintain their expression and function. Indeed, MYBL2 has been shown previously to transcriptionally regulate *POLQ* in lung adenocarcinoma [29]. However, we cannot rule out additional mechanisms of regulation of the DNA damage response by *MYBL2. MYBL2* is a multifunctional protein with a DNA binding domain at its N terminus, a central transactivation domain and protein interaction domain at its C terminus. Studies in somatic cells have shown an interaction between MYBL2 and the MRN complex at sites of DNA damage [26], which could also be the case for MYBL2 and Polθ/BRCA1/RAD51. Overall, we identify MYBL2 levels as a predictor of response to Polθ inhibitors in TNBC and observe that high MYBL2 levels are a vulnerability associated with increased drug sensitivity. Identifying patient groups which would most benefit from synthetic lethality approaches with Polθ inhibitor treatment is a critical step towards their translation into the clinic with a view to improving the outcomes for TNBC patients.

## Supporting information

within figures

## Acknowledgements

This work was funded by a Breast Cancer Now project grant (2019AugPR1320) awarded to PG. The authors wish to thank the staff at the Biomedical Services Unit at the University of Birmingham for their assistance with animal procedures. We would also like to thank Dr Mary Clarke and the staff at the University of Birmingham Flow Cytometry Unit for their support with cell sorting. We are very grateful to members of Clare Davies group including Dr Kelly Chiang and Dr Debashish Sahay for their help and guidance. pcDNA3.1(Hygro)myc-hPolQ-Flag was a gift from Agnel Sfeir (Addgene plasmid #73132; http://n2t.net/addgene:73132 ; RRID:Addgene_73132) [77] and RAD52-YFP [78] was a gift from Jiri Lukas.

## Author Contributions

Methodology RB, CD, PG, MS, Investigation, RB, AM, SA, CW, AS, BMB, KJ, Resources, RBC, GCMS, AMS, Writing - Original Draft RB, PG, Writing - Review & Editing RB, PG, AMS, MS, GCMS, RBC, CD, Funding acquisition PG, CD, Conceptualization PG, RB, Supervision PG, Project Administration PG,

## Declaration of Interests

GCMS is an employee and shareholder of ArtiosPharma Ltd and shareholder of AstraZeneca PLC. Experimental procedures

### Cell culture

MDA-MB-231, CAL-51 and 293T cells were cultured in Dulbecco’s modified Eagle’s medium (DMEM) (Gibco) supplemented with 10% foetal bovine serum (FBS) (Gibco), 50U/ml penicillin (Gibco) and 50µg/ml streptomycin (Gibco). shRNA mediated knockdown of MYBL2 was induced by addition of 1µg/ml doxycycline (Merck). MCF10A cells were cultured in DMEM/F-12 (Gibco) supplemented with 5% horse serum (Gibco), 20ng/ml EGF (Peprotech), 500ng/ml hydrocortisone (Merck), 100ng/ml cholera toxin (Merck), 10µg/ml insulin (Merck), 50U/ml penicillin (Gibco) and 50µg/ml streptomycin (Gibco).

### Generation of lentiviral plasmids containing MYBL2 full length or truncated versions of MYBL2

pHAGE-2-EF1a-Nterm truncated Mybl2-Linker-HA-SPA-EF1a-AmCyan, or pHAGE-2-EF1a-Cterm truncated Mybl2-Linker-HA-SPA-EF1a-AmCyan were generated from pHAGE-2-EF1a-Mybl2-Linker-HA-SPA-EF1a-AmCyan [25]. These plasmids contain mouse MYBL2 to avoid the silencing of human orthologs as human BC cell lines had been engineered with human MYBL2 shRNA. The N-terminal truncated version contains a 409 bp deletion between BmgBI (cuts at 50) and BstEII (cuts at 459) sites at the N-terminal of the protein. The C-terminal truncated version is 501bp truncated from the RsrII restriction enzyme site (cuts at 1599).

### Generation of stable cell lines

Stable cell lines were generated via lentiviral infection of mammary (MCF10A) or breast cancer (MDA-MB-231 and CAL-51) cells. For lentiviral production, 293T cells were transfected with pTRIPZ Inducible Lentiviral Negative shRNA Control (Horizon Discovery), pTRIPZ Inducible Lentiviral Human MYBL2 shRNA ORF (Horizon Discovery), pTRIPZ Inducible Lentiviral Human MYBL2 shRNA 3’UTR (Horizon Discovery), pHAGE-2-EF1a-AmCyan [25], pHAGE-2-EF1a-Mybl2-Linker-HA-SPA-EF1a-AmCyan [25], pHAGE-2-EF1a-N term truncated Mybl2-L-HA-SPA-EF1a-AmCyan or pHAGE-2-EF1a-C term truncated Mybl2-L-HA-SPA-EF1a-AmCyan, along with packaging vectors using trans-IT. Cells were cultured for 48 hours before harvesting and concentration of viral supernatant. Cells were transduced via centrifugation at 1500RPM for 2 hours at 32°C with viral supernatant containing 5µg/ml (MCF10A) or 8µg/ml (MDA-MB-231 and CAL-51) polybrene (Merck). Where appropriate cells were selected with 2µg/ml puromycin followed by sorting of RFP+ or AmCyan+ cells. For RFP expressing vectors the MOI was 10 and for AmCyan expressing vectors the MOI was 2.5. To generate MDA-MB-231 cells expressing luciferase for *in vivo* imaging, CMV-Luciferase (Bsd) lentiviral particles (amsbio) were transduced using the same method. The MOI used was 2 and cells were selected using 8µg/ml blasticidin. Luciferase activity was confirmed using the Luciferase Assay System (Promega) according to the manufacturer’s instructions.

### Mammosphere assay

Cell lines were detached from culture flasks using TrypLe (Gibco) and PDX tumour cells were dissociated using the human tumour dissociation kit (Miltenyi) according to the manufacturer’s instructions. Single cell suspensions were made by passing cells through a 25G needle and the number of live cells determined by tryphan blue exclusion. Cells were plated on pHEMA coated 6 well plates at a density of 3000 cells/well for CAL-51, 4500 cells/well for MDA-MB-231 and MCF10A, and 4750 cells/well for PDX cells. Cells were cultured for five (CAL-51, MDA-MB-231 and MCF10A) or seven days (PDXs) at 37°C. For MDA-MB-231, CAL-51 and PDXs, cells were cultured in DMEM/F12 phenol red free (Gibco) supplemented with B27 minus Vitamin A (Invitrogen), 20ng/ml EGF (Peprotech), and 50U/ml penicillin (Gibco) and 50µg/ml streptomycin (Gibco). For MCF10A, cells were cultured in DMEM/F12 phenol red free (Gibco) supplemented with B27 minus Vitamin A (Invitrogen), 20ng/ml EGF (Peprotech), 500ng/ml hydrocortisone (Merck), 100ng/ml cholera toxin (Merck), 10µg/ml insulin (Merck), 50U/ml penicillin (Gibco) and 50µg/ml streptomycin (Gibco). For drug treatments, 2.5µM ART558, 100nM AZ20, 2.5µM ART6043 or 2.5µM ART0380 were added on the day of seeding. Mammospheres >50µm were counted using an EVOS XL Core Imaging System (Thermofisher Scientific) and STEMgrid (STEMCELL Technologies). For secondary mammosphere formation, primary mammospheres were disaggregated into a single cell suspension using TrypLe (Gibco) followed by passing through a 25G needle. Cells were plated as for primary mammosphere formation and counted as previously. Counts were expressed as mammosphere forming unit (MFU).

### Isolation of anoikis resistant cells

Cells were plated at a density of 900 cells/cm^2^ on poly (2-hydroxyethyl methacrylate) (pHEMA) (Merck) coated dishes and cultured overnight at 37°C in the same medium used for the mammosphere assay. Anoikis resistant cells were purified using the Dead Cell Removal kit (Miltenyi) according to the manufacturer’s instructions.

### Isolation of CD24 ^low^/CD44^+^cells

CD24 /CD44 cells were purified sequentially using the CD24 Microbead kit (Miltenyi) followed by the CD44 Microbead kit (Miltenyi). Enrichment was determined by staining cells with anti-human CD24 Pe-Cy7 and anti-human CD44 APC-e780 (Invitrogen) and performing flow cytometry analysis using a BD LSR Fortessa X-20 (BD Biosciences).

### Western blotting

Cells were lysed in lysis buffer (20mM Tris pH 7.4/10mM EDTA/100mM NaCl/1% Triton-X 100) containing protease and phosphatase inhibitors. Protein extracts were clarified via centrifugation and protein concentration determined using a Bradford reagent (Bio-Rad). 50µg protein extract was separated by SDS-PAGE and transferred to a PVDF membrane via a wet transfer system (Bio-Rad). Membranes were blocked for 1 hour at RT with 5% dried milk/TBST, incubated with primary antibodies diluted in 5% dried milk/TBST overnight at 4°C and then HRP-conjugated secondary antibodies diluted in 5% dried milk/TBST for 1 hour at RT. Protein signals were visualised using ECL detection reagents (GE Healthcare).

### ALDEFLUOR assay

ALDEFLUOR assay (STEMCELL Technologies) was carried out according to the manufacturer’s instructions by incubating the ALDEFLUOR dye with 1×10^5^ cells per sample for 30 minutes at 37°C.

### RNA extraction, cDNA and qPCR

RNA was extracted using the RNeasy mini or micro kits (Qiagen) according to the manufacturer’s instructions, including the removal of contaminating genomic DNA through treatment of samples with RNAse-free DNAse (Qiagen). Reverse transcription to generate cDNA was performed using the SuperScript VILO cDNA Synthesis Kit (ThermoFisher Scientific) according to the manufacturer’s instructions. qPCR was carried out using TaqMan probes (ThermoFisher Scientific) and TaqMan Gene Expression Master mix (Applied Biosystems) or KiCqStart SYBR Green Primers (Merck) and PowerUp SYBR Green Master Mix (Applied Biosystems), all according to the manufacturer’s instructions. Beta-2 microglobulin was used as a control to standardise samples.

### Luminescence-based DNA repair assays

DNA repair reporter substrates (Artios Pharma Limited) have been engineered to express a functional NanoLuciferase reporter protein only when HR, NHEJ, SSA or TMEJ have been correctly performed [49, 51, 79]. Plasmids were cotransfected with a Firefly vector (pGL4.50[luc2/CMV/Hygro], Promega) to provide a control for transfection efficiency and cell density. Anoikis resistant CAL-51, MDA-MB-231 or MCF10A cells were seeded into white 96 well tissue culture plates (ThermoFisher Scientific) at a density of 1×10^4^ (MDA-MB-231 and CAL-51) or 2x10^4^ (MCF10A) cells/well for 24 hours at 37°C. Each well was transfected with 5µl optimem, 0.25µl Lipofectamine 3000, 0.2µl P3000 reagent (ThermoFisher Scientific) and the appropriate plasmid combinations according to the manufacturer’s instructions. These included (1) 80ng HR reporter + 25ng Firefly, (2) 20ng NHEJ reporter + 25ng Firefly, (3) 40ng SSA reporter + 25ng Firefly, or (4) 20ng TMEJ reporter + 80ng Firefly. After 24 hours, Firefly and NanoLuciferase levels were determined using the Nano-Glo Dual-Luciferase Reporter system (Promega) according to the manufacturer’s instructions. Luminescence was measured using a GloMax Explorer plate reader (Promega) using the luminescence setting using a 1 second integration time. Nanoluciferase signal was normalised to Firefly signal and results are expressed through comparison to the appropriate experimental control (shCTRL or AmCyan).

### Colony survival assay

Bulk or anoikis resistant MCF10A, CAL-51 or MDA-MB-231 cells were plated at low density and exposed to different concentrations of Olaparib (Selleckchem), ART558 (Artios Pharma Limited) or AZ20 (Selleckchem). For experiments using doxycycline inducible shRNA-mediated knockdown medium was refreshed every 2-3 days. After 10 days, colonies were fixed and stained with 0.5% crystal violet (Merck) in ddH_2_O. Colony counts are expressed as percentage survival when compared to cells treated with DMSO.

### BrdU assay

Cells were cultured in the presence of 25µM 5-bromo-2-deoxyuridine (BrdU) (Merck) and harvested for fixation in 75% ethanol for 30 minutes at 4°C. Cells were rehydrated in PBS for 20 minutes at RT and denatured using 2N HCl for 20 minutes at RT. Cells were washed twice in PBS and twice in 5% FBS/0.1% NaN3/0.1% Triton-X 100/PBS before incubation for 50 minutes at RT with FITC Mouse Anti-BrdU or FITC Mouse IgG1 κ Isotype Control (BD Biosciences). Cells were washed twice with PBS and then flow cytometry performed using a BD LSR Fortessa X-20 (BD Biosciences).

### DNA fibre analysis

S1 DNA fibre analysis was carried out as described previously with minor modifications [80]. Anoikis resistant cells were cultured in the presence of 25µM chlorodeoxyuridine (CIdU) (Merck) for 20 minutes at 37°C. Cells were washed two times in PBS and cultured in the presence of 250µM iododeoxyuridine (IdU) (Merck) for 60 minutes at 37°C. Cells were washed two times in PBS and permeabilised using CSK100 buffer (100mM NaCl/10mM HEPES/3mM MgCl_2_/300mM sucrose/0.5% Triton-X 100) for 10 minutes at RT. Nuclei were incubated with S1 buffer -/+ 20U/ml S1 nuclease (ThermoFisher Scientific) for 20 minutes at 37°C. Cells were removed from wells using a cell scraper, centrifuged and resuspended in PBS at 1500 cells/µl. Fibre spreading and immunostaining were carried out as described previously [81]. Images were captured using a Leica DM6000B microscope at x40 magnification and fibre lengths measured using LAS X software (Leica).

### Immunofluorescence

Anoikis resistant MCF10A, CAL-51 and MDA-MB-231 cells were grown on glass coverslips. Alternatively, mammospheres generated from MCF10A, CAL-51 and MDA-MB-231 cells were disaggregated into a single cell suspension using TrypLe (Gibco) followed by passing through a 25G needle. Single cell suspensions were cytospun onto glass coverslips. Cells were irradiated with 3Gy of ionising radiation or treated with 10µM cisplatin to induce DNA damage. For detection of Rad52 and Polθ foci, anoikis resistant cells were transfected with 2.5µg pcDNA 3.1(Hygro) myc-hPolQ-Flag plasmid [77] or RAD52-YFP [78] and incubated for 24 hours prior to induction of DNA damage. At indicated time points, cells were permeabilised with nuclear extraction buffer (20mM NaCl/3mM MgCl_2_/300mM sucrose/10mM PIPES/0.5 % Triton X-100, pH 6.8) for 5 minutes on ice, followed by fixation in 4% PFA for 10 minutes at RT. Cells were washed three times in PBS and blocked for 1 hour at RT with 10% FBS/PBS. Antibody staining was carried out as described previously [82]. Images were captured using a Leica DM6000B microscope at 40x magnification and foci numbers counted using LAS X software (Leica).

### Immunohistochemistry

Tumours were fixed with 10% neutralised buffered formalin (NBF) (Merck) and stored at 4°C before being embedded in paraffin. Sections were stained with haematoxylin (Merck) and eosin (Merck) according to the manufacturer’s instructions by the Human Biomaterials Resource Centre at the University of Birmingham.

### Xenograft studies and *in vivo* imaging

All animal experiments were conducted in accordance with the United Kingdom Home Office Regulations. 7–9-week-old female NSG (NOD.Cg-Prkdcscid Il2rgtm1Wjl/SzJ) mice received a subcutaneous injection of the appropriate number of MDA-MB-231 shCTRL or shMYBL2 cells at a 1:1 ratio with Matrigel (Corning) into the left flank. Prior to injection cells had been cultured in the presence of 1µg/ml doxycycline and animals were maintained on dox diet (Envigo) throughout the experiment to induce shRNA-mediated gene knockdown. Tumour growth was monitored using caliper measurements and IVIS imaging. 15 minutes prior to IVIS imaging, animals received an intraperitoneal injection of 150µg/g luciferin (Promega) and images were captured on an IVIS Spectrum Imaging System. At the experimental end point, tumours were harvested, photographed and the weight recorded. Tumours pieces were stored in NBF to be used for IHC or snap frozen and stored at -80°C for protein extraction. Remaining tumour pieces were dissociated into single cells using the Miltenyi Tumour Dissociation Kit (Miltenyi) and used for mammosphere assays.

### *In vivo* implantation of patient-derived xenograft models

All animal experiments were conducted in accordance with the United Kingdom Home Office Regulations. 7–9-week-old female NSG (NOD.Cg-Prkdcscid Il2rgtm1Wjl/SzJ) mice were implanted with two ∼3mm^3^ PDX tumour fragments [83] coated in Matrigel (Corning). Fragments were implanted into a subcutaneous skin pocket, on the left side and adjacent to the fourth mammary fat pad. Tumour growth was monitored using caliper measurements. At the experimental end point, tumours were harvested, photographed and the weight recorded. Some tumour pieces were frozen and the rest were dissociated into single cells using the Miltenyi human tumour dissociation kit (Miltenyi) and used for RNA extraction, protein extraction and mammosphere assays.

### Chromatin Immunoprecipitation

Chromatin immunoprecipitation was carried out as described previously [84]. Briefly, bulk or anoikis resistant MDA-MB-231 cells were crosslinked with 1% formaldehyde and neutralized using 0.125M glycine. Cells were lysed, nuclei isolated and sonicated to shear DNA to 300-1000bp fragments. Samples were pre-cleared with mouse immunoglobulins (Santa Cruz Biotechnology) and chromatin was co-immunoprecipitated with 5µg of anti-MYBL2 antibody (Santa Cruz Biotechnology) or anti-mouse immunoglobulin antibody (Santa Cruz Biotechnology) overnight at 4°C while rotating. Protein-DNA complexes were incubated with Protein G magnetic beads (Pierce) for 3 hours at 4°C while rotating. Samples were washed and complexes eluted from the beads, crosslinks were reversed and proteinase K treatment performed. DNA was purified using a QIAquick PCR Purification Kit (Qiagen) and quantified using a Nano Drop 2000 (ThermoFisher Scientific). ChIP efficiency was measured using qPCR. Samples were compared to input chromatin and quantified through comparison to immunoglobulin samples.

### Analysis of The Cancer Genome Atlas

TCGA Firehose Legacy (Breast Invasive Carcinoma) data was obtained from cBioPortal [58-60]. Samples were profiled for MYBL2 mRNA expression z-scores relative to all samples with a z-score threshold of 1.0. Samples were split into low and high MYBL2 expression based on 1 standard deviation below and above the mean mRNA expression level. These criteria allowed analysis to be performed on 193 MYBL2low samples and 190 MYBL2high samples. The Comparison/Survival analysis function was used to extract the data.

### Analysis of the Cancer Cell Line Encyclopaedia

Cancer Cell Line Encyclopaedia data was obtained through the DepMap portal (https://depmap.org/portal) [85]. The Batch corrected Expression Public 24Q2 dataset was analysed using Pearson Correlation for MYBL2 expression in breast cancer cell lines and plotted against expression of genes of interest.

## Statistical analysis

Statistical differences for foci numbers, DNA fibre length and mammosphere size were calculated using the Mann-Whitney rank sum test. For all other experimental data, statistical differences were determined using the Student’s t-test. Statistical tests were carried out using GraphPad Prism Version 10.3.0 (GraphPad Software) through comparison to control-treated samples. * p=<0.05, *** p=<0.01 and *** p=<0.001.

