## Supplementary material for "MYBL2 maintains stemness and promotes theta-mediated end joining in triple negative breast cancer": within figures

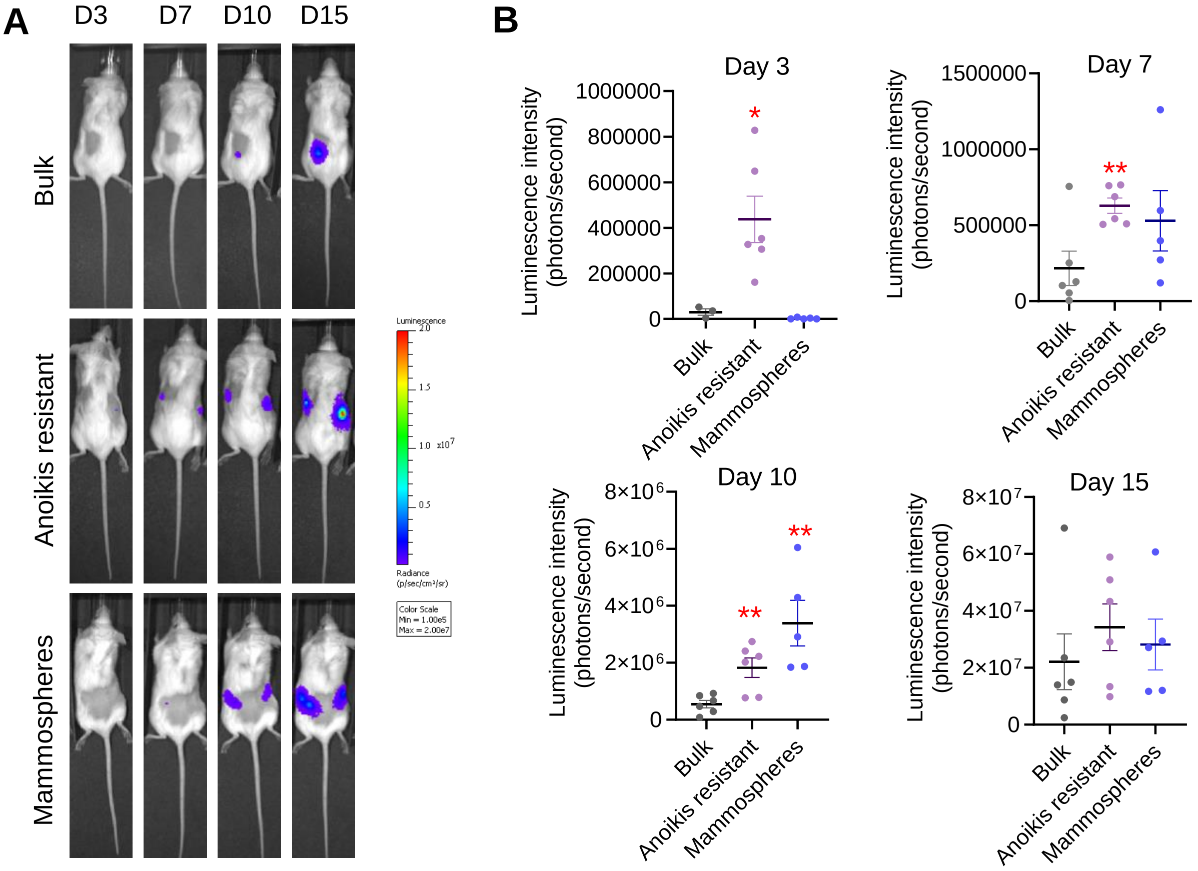


**Supp Figure 1. Triple negative breast cancer stem/progenitor cells purified using anoikis resistance or through mammosphere assays have increased tumour initiation capacity *in vivo* compared to the bulk cell population.** Mammospheres were generated or anoikis resistant cells isolated from the bulk population of MDA-MB-231 cells. Cells were injected into the flank of NSG mice. (A) Representative IVIS images over time of animals injected with bulk, anoikis resistant or mammosphere derived MDA-MB-231 cells. (B) Quantification of IVIS images on different days post injection of bulk, anoikis resistant or mammosphere derived MDA-MB-231 cells. Plots represent data from five-six independent tumours are shown as mean±SEM. *p ≤0.05, **p ≤0.01 and ***p ≤0.001 and obtained from a two-tailed t test.

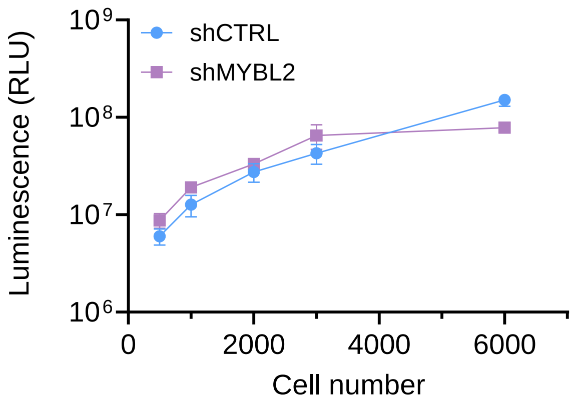


**Supp Figure 2. shCTRL and shMYBL2 MDA-MB-231 isogenic cells display similar levels of luminescence *in vitro*.** shCTRL and shMYBL2 MDA-MB-231 cells constitutively expressing the luciferase gene were counted, harvested and lysed. Different numbers of cells were plated out and luminescence was measured following addition of substrate. Plot represents data from one independent experiment measuring luminescence from three technical replicates for each cell number and shows the mean±SEM.

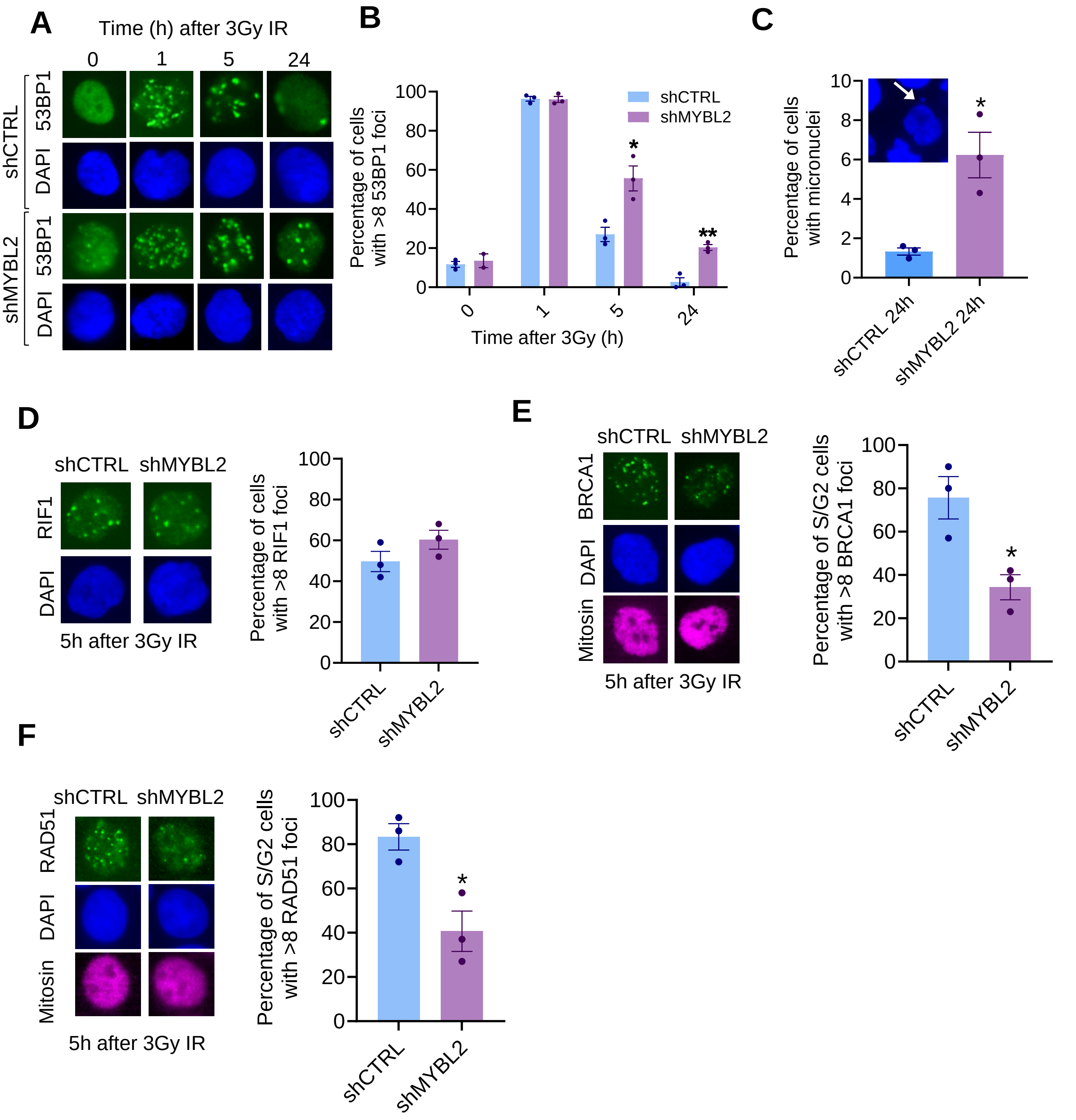


**Supp Figure 3. MYBL2 supports DNA double strand break repair in cell populations enriched for triple negative breast cancer stem/progenitor cell activity derived from bulk CAL-51 cells.** (A+B) Mammosphere cultures of CAL-51 cells expressing pTRIPZ Inducible Lentiviral shRNA control (shCTRL) or MYBL2 (shMYBL2) were incubated with doxycycline for 5 days, exposed to ionizing radiation (IR) and immunostained with antibodies to 53BP1 at different time points. Representative microscopy images are shown (A) and quantification of 53BP1+ cells (B). (C) Mammosphere cultures were treated with doxycycline as in (A+B), exposed to IR and micronuclei formation assessed 24h later. (D-F) Mammosphere cultures were treated with doxycycline as in (A+B), exposed to IR and immunostained with antibodies to RIF1 (D), BRCA1 and mitosin (E) or RAD51 and mitosin (F). For each panel representative microscopy images and foci quantification are shown. Plots represent data from at least three independent experiments and are shown as mean±SEM. *p ≤0.05 and **p ≤0.01 and obtained from a two-tailed t test.

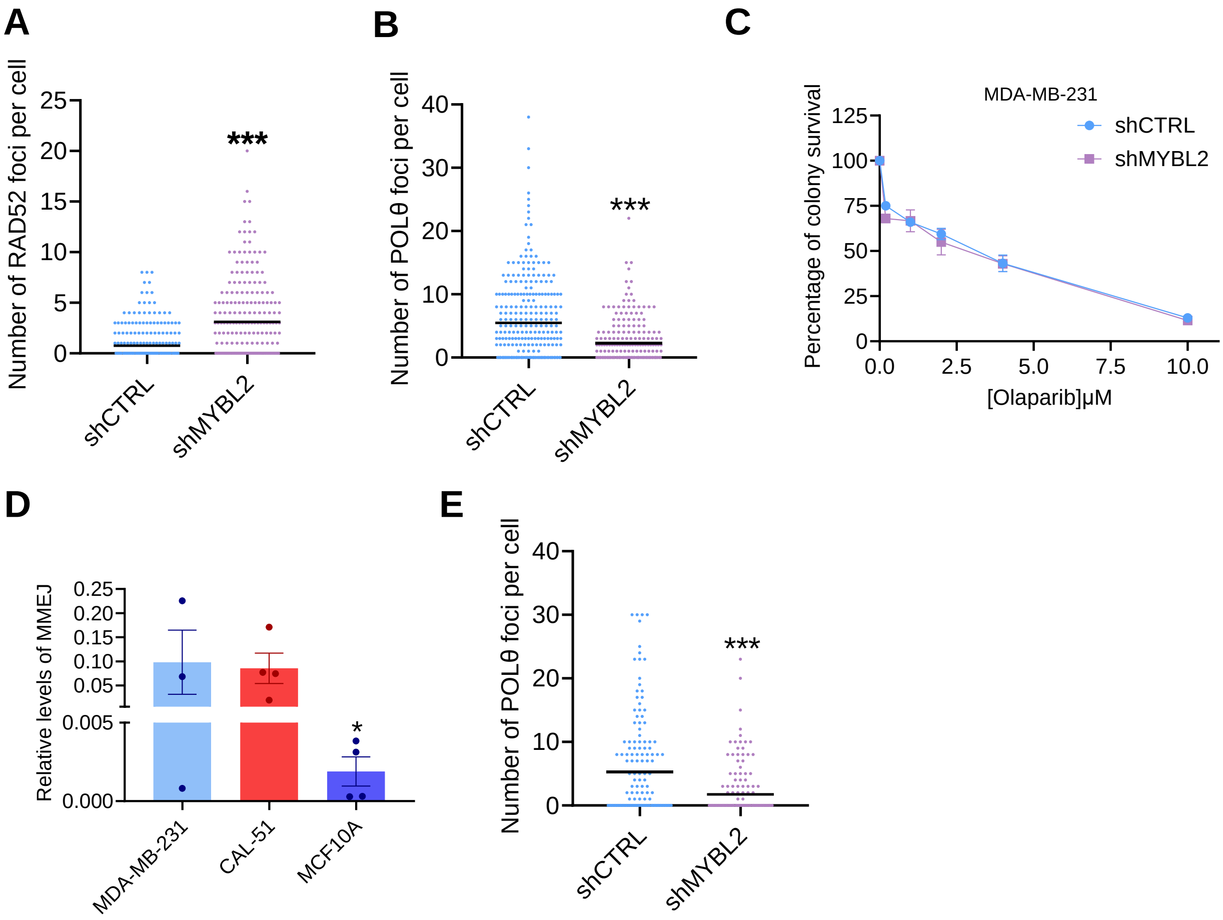


**Supp Figure 4. Cell populations enriched for breast cancer stem/progenitor cell activity display differential recruitment of DNA repair proteins depending on levels of MYBL2.** (A+B) Anoikis resistant cells were isolated from bulk MDA-MB-231 cultures expressing pTRIPZ Inducible Lentiviral shRNA control (shCTRL) or MYBL2 (shMYBL2). Cells were transfected with RAD52-YFP (A) or Polθ-FLAG (B) plasmids, exposed to ionizing radiation (IR) and imunostained with antibodies to mitosin (A) or FLAG and mitosin (B). Plots show the number of foci present in each cell counted. (C) Anoikis resistant cells were isolated from bulk shCTRL or shMYBL2 MDA-MB-231 cultures. Cells were treated with olaparib and doxycycline and left for 10 days to form colonies. Colonies were stained with methylene blue and counted. (D) Anoikis resistant MDA-MB-231, CAL-51 or MCF10A cells were transfected with an TMEJ NanoLuciferase report plasmid. Cells performing DNA repair via TMEJ were quantified via measurement of luminescence. (E) Anoikis resistant shCTRL and shMYBL2 cells isolated from bulk MDA-MB-231 cultures were transfected with Polθ-FLAG plasmid, treated with cisplatin and immunostained with antibodies to FLAG and mitosin. Plot shows the number of foci present in each cell counted. All plots represent data from at least three independent experiments and are shown as mean±SEM. *p ≤0.05 and ***p ≤0.001 obtained from a Mann-Whitney test (A-B+E) or from a two-tailed t test (D).

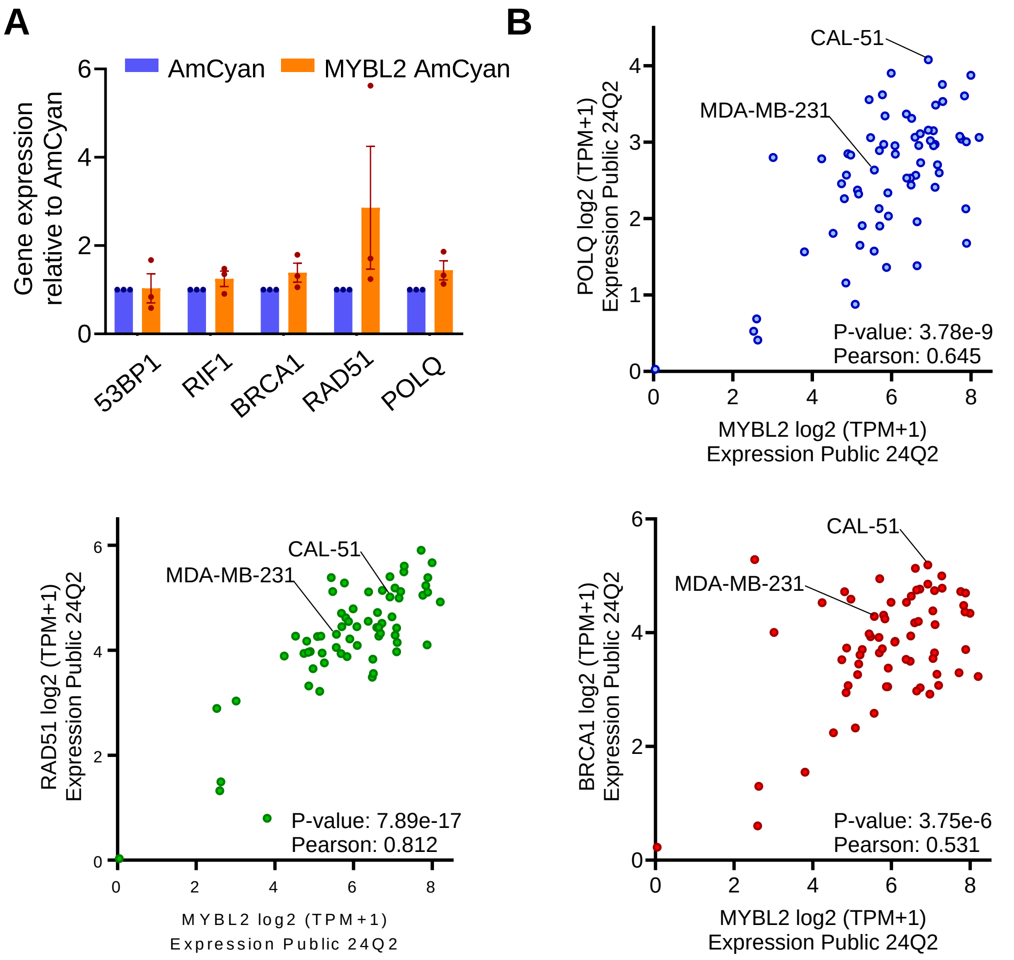


**Supp Figure 5. Expression of DNA repair genes in normal mammary stem cells and the bulk cell population in breast cancer cell lines depending on levels of MYBL2.** (A) Mammospheres were generated from MCF10A cells expressing pHAGE-2-EF1a-L-HA-SPA-EF1a-AmCyan (AmCyan) or pHAGE-2-EF1a-Mybl2-L-HA-SPA-EF1a-AmCyan (MYBL2 AmCyan) and expression of the indicated genes measured by qPCR. Plot represents data from three independent experiments and are shown as mean±SEM. (B) Correlation between MYBL2 mRNA expression and mRNA expression of DNA repair genes in different breast cancer cell lines. Data extracted from the Cancer Cell Line Encyclopedia. P-values shown were obtained using Pearson Correlation.
